# Gaining confidence in inferred networks

**DOI:** 10.1101/2020.09.19.304980

**Authors:** Léo P.M. Diaz, Michael P.H. Stumpf

## Abstract

Network inference is a notoriously challenging problem. Inferred networks are associated with high uncertainty and likely riddled with false positive and false negative interactions. Especially for biological networks we do not have good ways of judging the performance of inference methods against real networks, and instead we often rely solely on the performance against simulated data. Gaining confidence in networks inferred from real data nevertheless thus requires establishing reliable validation methods. Here, we argue that the expectation of mixing patterns in biological networks such as gene regulatory networks offers a reasonable starting point: interactions are more likely to occur between nodes with similar biological functions. We can quantify this behaviour using the assortativity coefficient, and here we show that the resulting heuristic, *functional assortativity*, offers a reliable and informative route for comparing different inference algorithms.

## 1 Introduction

Network inference is the process by which we aim to learn the structure of networks from data [1, 2]. The networks that we are particularly interested in are those that capture molecular signalling and regulatory processes. However, such interactions occurring inside cells are often hard to observe, and statistical dependencies between indirect observations are used as a proxy to infer real interactions in the processes of interest. That way, dependency in patterns of gene expression may be taken as a reflection of real interactions between e.g. the genes or their products, but such relationships are particularly difficult to infer indirectly [3].

There is a vast literature on developing approaches for network inference (reviewed partially in [1, 4, 5, 2]). The panoply of methods includes: correlation and partial correlation measures; Bayesian network algorithms; information-theoretical dependency measures; regression approaches; methods adapted from dynamical systems theory; general machine learning approaches, including different flavours of deep neural networks; and hybrid methods that incorporate a panel of different estimation procedures. Each method comes with its own set of assumptions and limitations, and these may not always be made explicit.

Assessing the strengths and weaknesses of different methods, and comparing their performance has been fraught with difficulties, such as the high computational cost of many network inference methods, which has often prohibited extensive analysis [2]. More importantly, however, is the scarcity of suitable test datasets, with large, exhaustively validated networks of real biological systems remaining largely elusive. The DREAM initiative is an ongoing effort aiming to remedy this lack of ground truth to use as reference by providing solid *in silico* test cases for which we can precisely evaluate and compare the performance of different statistical approaches, including network inference methods, which were the focus of the DREAM 4 challenges for instance [4]. Other studies have followed up on this to provide similar assessments of network inference methods [5].

Yet, conclusions drawn from such efforts also come with limitations. Worryingly, these may be easily overlooked, often as a consequence of the design setup of the challenges themselves, presented as contests where inference algorithms are ranked from best to worst according to their performance. Such rankings in absolute terms are quick to discard the specific context in which an algorithm was tested as *in silico* tests may have implicit or explicit biases for a particular set of approaches over others. Therefore such rankings are only valid in the specific, highly controlled setting of the corresponding inference challenge [6].

In some instances certain algorithms have become *de facto* standards, either because they arrived early on the scene or because of their fast or easy implementation; and often less emphasis is put on assessing their accuracy, with the quality of their predictions rarely being evaluated explicitly post publication. Sometimes, and this is demonstrably not appropriate, inferred networks have even been analysed as if they were reliable representations of biological reality.

Clearly the situation is far from satisfactory: (i) there is clear need for better models of biological systems, including networks, which can form the basis for more detailed mechanistic and predictive models; (ii) *in silico* methods could be a cheaper and attractive alternative to many experimental assays, provided their limitations are made explicit; (iii) apart from sanitised simulated data there is typically very little to go on for a meaningful evaluation of an algorithm’s performance.

Here we introduce and discuss a heuristic that allows us to quantify relatively the confidence we should have in proposed biological networks, such as those emerging from network inference. Heuristics of this type – and we shall revisit and stress this point below – offer primarily a sanity check: if the inferred network scores very poorly, we should probably resist from analysing it further. The heuristics are not meant to replace experimental or statistical (in)validation [7, 8] rather they aim to put on a quantitative basis what is frequently done by visual inspection.

Below we first outline network inference and the plausibility of inferred networks; we then illustrate how *network assortativity* [9, 10] allows us to compare and rank different network inference algorithms; we then outline how this approach can be employed in practice, before concluding with a discussion on difficulties in the process of network inference.

## 2 Assessing the plausibility of inferred networks

A network is represented by the ordered pair

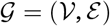

where *V* denotes the set of nodes or vertices *ν* = *{v*_1_, *v*_2_, …, *v*_*N*_ *}*, and *ε* = *{e*_1_, *e*_2_, …, *e*_*M*_ *}*, set of links or edges. While *V* is typically known, *E* only is in a few instances, and, arguably, exceedingly rarely in biology; instead we rely on statistical methods to infer the presence or absence of edges between pairs of nodes *v*_*i*_, *v*_*j*_, *∈,V i, j* = 1, …, *N*. We will not distinguish between directed and undirected networks as our discussion is applicable to both with only minor modification.

Network inference algorithms typically score edges [1, 2], and this score, here denoted by *ξ*_*ij*_, represents the relative weight in favour of an edge existing between nodes *v*_*i*_ and *v*_*j*_. We shall often write *ξ*(*q*), to denote the *q*th highest score (we ignore possible ties, which can be straightforwardly resolved by ordering them randomly), and understand that this refers to the score of the corresponding edge. Network inference is thus based on a process by which a pair of nodes is assigned a real value,

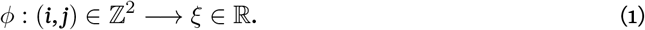

In fact the function *ϕ* really takes states, *η*_*i*_ and *η*_*j*_, associated with nodes, *i* and *j*, to determine the scores, *ξ*,

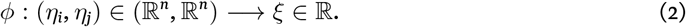

For a set of *l* network inference methods,

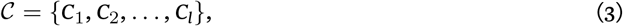

which will result in inferred sets of edges, *ε*_1_, *ε*_2_, …, *ε*_*l*_, we want to assess the relative merit of these *candidate* inferred networks, which are, within the constraints of the methodology, the best available representation of the real network of interest.

### 2.1 Properties of biological networks

Any real biological network (we note that there are limitations to networks as representations of real-world biological systems) is expected to have certain properties, which include

1. Specificity: interactions will be more likely between nodes that have certain functionality (e.g. belong to the same functional class; or belong to different functional classes that have a high probability of interacting – here *Gene Ontology* annotations can serve as a proxy for, or best guess of, functionality).
2. Modurality: groups of nodes will form tightly interacting modules with pronounced clique structure to fulfil their biological function; modules are expected to be enriched for nodes that have similar or related functions.
3. Connectedness: the true network will connect all nodes (this is not necessarily the case for incomplete data [11]).
4. Robustness: gross structural features, and thus thefunction of thenetwork, should be robust against the removal of individual nodes.
5. Hierarchy: some nodes will have more prominent network positions (degree, centrality) and may orchestrate module and modular dynamics.
6. Balance: a real network should have a structure that reflects function and functional importance [12, 13]. For similar importance we can expect similar levels of network organisation, robustness, and modularity across the whole network [14].

None of these points should be contentious if we accept (with the usual *caveats*) the functional relevance of biological networks. These points may contradict some simplistic network models [15], but, as has been argued, and indeed demonstrated, elsewhere, the structure of real biological networks is much more nuanced and “scale-rich” than simple models might have suggested [12, 16, 14].

Point 1, in particular (and to a lesser extent also point 2), allows us to develop quantitative criteria against which proposed networks (here we are predominantly concerned with inferred networks) can be evaluated. Points 3 and 4 reflect on network properties that go beyond local interactions, which may nevertheless help tocompare the performance of differentnetwork inference methods [3, 11]. Forpoints 5 and 6 we may also be able to develop testing procedures, but these would have to start more explicitly from the top-down: coarsegraining and renormalisation methods may offer some potential routes [22].

One important distinction needs to be made regarding the types of node properties we may want to compare in points 1 and 2. They can be categorical or structural: among the former we include biological annotations [23]; among the latter network properties of nodes [9, 10]. For the former we can assume a null-model of independence. For the latter we can only assume conditional independence (conditional on aspects of network structure) which makes testing more complicated [23].

### 2.2 Quantifying aspects of network organisation through assortativity

Mixing patternsrefertotheoverall networkorganisation arising through attachmentof nodes toothernodes with similar properties, and for pairwise comparisons we can use the *assortativity coefficient* [9, 10] to quantify this behaviour. This assumes that we can assign each node to a set of *q* properties, *K* = *{κ*_1_, *κ*_2_, …, *κ*_*q*_ *}*; here *κ*_*q*_ may represent “unknown”. Crucially, the properties *κ*_*i*_, *i* = 1, …, *q* must be different from the measurements or states, *η*_*j*_, *j* = 1, …, *u* that were used for inferring the network [23].

The number of nodes with annotation *κ*_*i*_ is denoted by *ν*_*i*_. We then define a matrix, *A*, where the entries, *a*_*ij*_, are the number of edges connecting nodes with annotation *i* with those with annotation *j*. The assortativity coefficient [10], *r*, then is given by

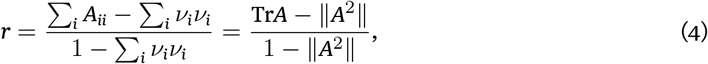

where the second equality results straightforwardly from conventional properties of matrix representations of networks.

The assortativity coefficient quantifies mixing patterns: confined to the range *–* 1 *≤ r ≤* 1, a network is said to be assortative when *r >* 0 (where nodes tend to be connected to nodes with similar properties), and disassortative otherwise [9].

The assortativity coefficient was originally calculated using node degree as a basis to compare node similarity, yielding *degree assortativity* [9]. However, in addition to node degree, any other node annotation may be used.

Functional network modules play a crucial part in cellular processes [24, 25, 26, 27], and inferred networks should reflect this organisation. Quantifying network assortativity with respect to functional annotations of nodes then allows us to draw from both points 1 and 2 in Sec. 2.1, (functional) specificity and modularity: assortativity can be used as a heuristic to quantify the explicit assumption of mixing patterns by biological function.

Experimental evidence supports the importance of functional modules in biological networks, from observations in *Saccharomyces cerevisiae* of preferential interaction between functionally related genes [28, 23, 29] that cluster at the level of cellular process [17] into functional modules with more connections within modules than between than expected in random networks [30], to the identification of groups of gene (“dynamical modules”) coherently implementing biological functions in the *Drosophila melanogaster* gap gene network [27]. The clustering of genes within biological process supports the assumption of functional modules, i.e. mixing patterns with respect to biological function.

As we have argued, this behaviour is quantified by the assortativity coefficient: under this assumption, we expect biological networks to exhibit assortative mixing with respect to biological function, and a higher coefficient indicates more support in favour of a given network. We refer to this heuristic as *functional assortativity*, which is a function of node annotations corresponding to biological function. This proxy measure for quantifying the *plausibility* of inferred networks presents the advantage to hold regardless of the inference methodology and thus allows to rigorously compare inference algorithms.

## 3 Measuring confidence in inferred networks

Below we outline the inference methods used, before discussing their respective candidate networks in light of the assortativity coefficients.

### 3.1 Inference algorithms considered

We compare the performance of seven inference algorithms and use these to illustrate the behaviour of the assortativity coefficient. We use two correlation-based approaches – linear correlation (LC) and rank correlation (RC) coefficients – and an information-theoretic approach – based on the mutual information (MI) – as baseline predictions because of their popularity and ease of use (e.g. [31]); to these we add three other information-theoretic approaches – context likelihood of relatedness (CLR) [20], proportional unique contribution (PUC) [3], and partial information decomposition and context (PIDC) [32, 3] – and a regression-based algorithm – GENIE3 [21], ran here with default settings – see Table 1 for more detailed descriptions of each. The focus on information-theoretic approaches stems from the ability of mutual information to capture non-linearities [19, 33], which is of obvious importance in a biological context.

**Table 1:** Description of inference algorithms compared.

| algorithm | description | reference |
| --- | --- | --- |
| linear correlation | Measures the linear correlation between a pair of random variables. | [17] |
| rank correlation | Measures the rank correlation between a pair of random variables. | [18] |
| MI | Measures dependency between variables using the mutual information, that is the sum of the entropy of the variables minus their joint entropy; it represents the amount of information about one variable when another variable is known. | [19] |
| CLR | Based on the value of the MI between pairs of variables in the context of MI scores for each possible combination of variable pairs. This approach is referred to as *network context* and amounts to calculating the likelihood of each MI score conditional on the overall score distribution. | [20] |
| PUC | Based on the mean unique information between variable pairs that accounts for their MI, as calculated via the partial information for each possible variables triplet for a given pair. | [3] |
| PIDC | Builds on the PUC approach by taking the network context into account in a similar way that CLR does i.e. by considering the overall distribution of PUC values. | [3] |
| GENIE3 | Creates as many regression problems as the number of input genes, then uses random forests to infer edges and their nature (genes are considered putative TFs if setting them as nodes on the trees reduces the variance of the predicted output). | [21] |

We choose to focus on undirected networks; that way, assumptions about putative regulatory relationships are kept minimal and each edge can be treated as a falsifiable hypothesis. GENIE3 [21] produces directed networks, which we make undirected to allow comparison; we did this by retaining only the first occurrence of each edge in either direction (meaning that each edge in the undirected network is ranked according to the position of the most likely interaction in the directed network).

We illustrate the methods by applying these inference algorithms to a single cell dataset of mouse embryonic stem cells, where gene expression is measured over seven days as cells differentiate into neurons [34]. Each gene is manually annotated with one 12 classes of biological functions (mesoderm, primitive endoderm, endoderm, neuroectoderm, trophoectoderm, naive pluripotency, primed pluripotency, core pluripotency, loading control, cell cycle, chromatin modulator, and signalling), which allows us to measure functional assortativity as described above.

### 3.2 Functional assortativity coefficient

We plot the functional assortativity coefficient (FAC) as a function of the number of candidate edges included in the networks resulting from the different methods in Fig. 1. By necessity this is either 1 or −1 depending on whether the first edge is between nodes with the same or with different annotations. Both can be biologically reasonable: a different annotation could for example result when one nodes is annotated as “primed pluripotency” and the other node as “signalling”, as is the case for the top-rated edge resulting from PIDC (which connects CLDN6 and IGF2); this is a biologically plausible, and in line with known relationships in several organisms. The same annotation of both nodes, is indicative of functional relationship as outlined above; “core pluripotency”, for example, is shared by FGF4 and POU5F1/OCT4, the top-ranked edge for CLR, PUC, MI, and RC, and the 8th highest ranked edge for PIDC; this is a well-documented interaction playing a central role in stem cell differentiation [35, 36, 37].

**Figure 1:**
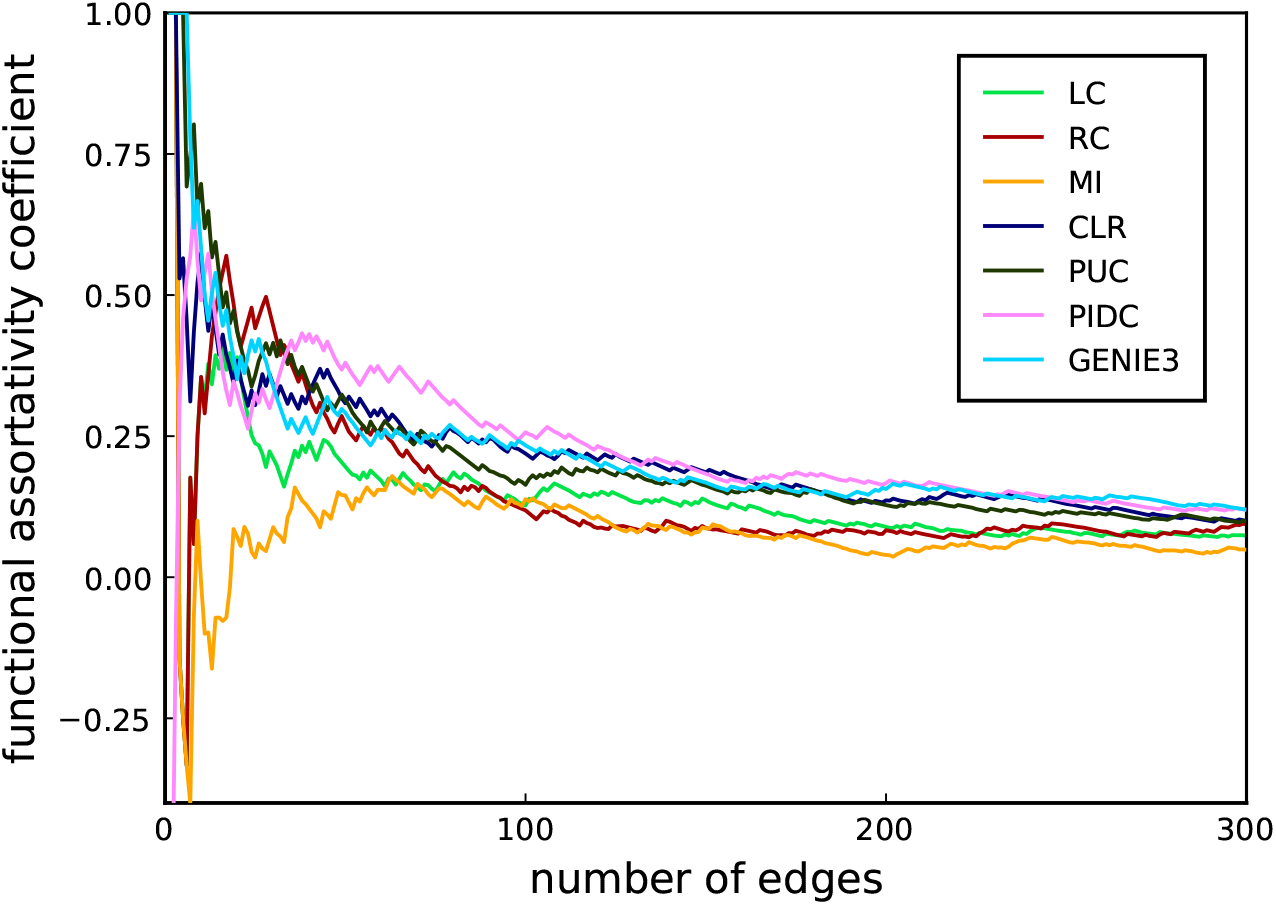
Evolution of the FAC as a function of the number of edges in a relevance network where edges are introduced in the order implied by their score.

It is possible to work down the whole list of interactions and seek explicit confirmation for each scored interaction. This could be subject to investigator bias, and the rationale for using the assortativity coefficient is to make this process automated and, conditional on the available network and annotation data, unbiased. So while a realistic network will have – even for high-quality and nuanced annotations – a proportion of cross-category edges, a majority of within-category edges is expected.

The three more advanced information-theoretic inference methods, PIDC, CLR and PUC, display the highest FAC at a given edge number (Fig. 1). The assortativity coefficient for every inference method eventually decreases into the background noise as the networks becomes a completely connected graph, as expected. For each inference method we observe a maximum in the FAC for low to moderate values of the number of edges included in the network (roughly between 50 and 150). For every value of the number of edges included in the network, the FACs, for networks inferred with PIDC, CLR and PUC are generally higher than the FACs obtained for the other networks.

This demonstrates that these algorithms result in inferred networks that have a higher number of interactions among functionally related nodes, compared to correlation or mutual information. As this is in line with biological knowledge and intuition, we would put more trust into networks inferred with e.g. CLR, PUC or PIDC than networks inferred by other means.

### 3.3 Discrepancies in inference algorithms predictions

The different inference algorithms *l*, yield different sets of inferred edges, *ε*_*l*_, as is obvious in the overlap patterns of the Venn diagrams shown in Fig. 2: while most edges are shared across inference algorithms, each method infers a set of interactions that no other methods pick up. While this is already known and is consistent with observations of discrepancies in widely used between inference methods for single-cell data [38], it also further highlights the need for developing better ways to assess our confidence in inferred networks, especially in the absence of ground truths.

**Figure 2:**
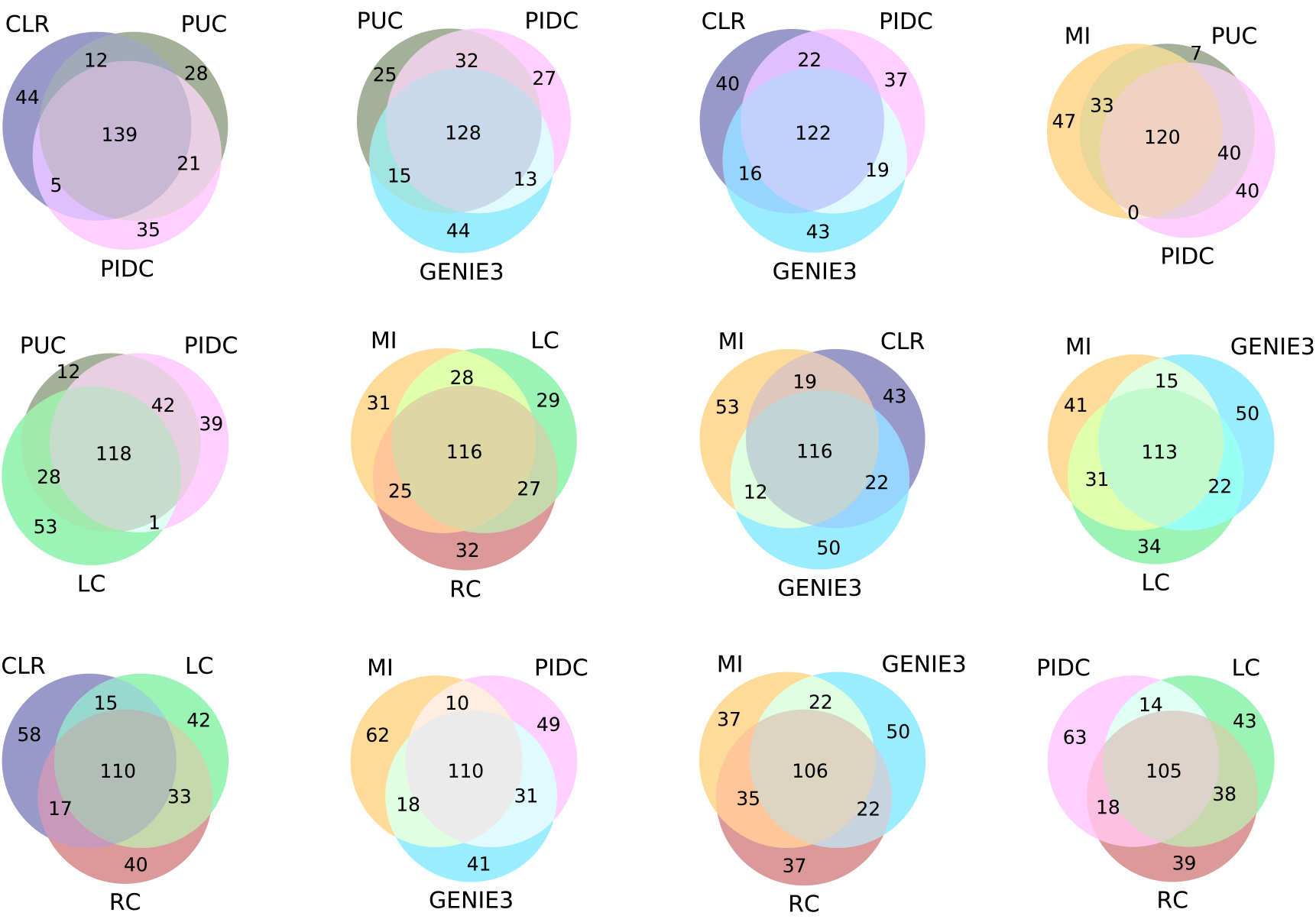
A selection of Venn diagrams showing patterns of overlap between three given inference methods for relevance networks with 200 edges. Overlaps are according to the number of edges shared between the given inference methods. Large overlap can mean that the different methods detect the same signal, which does not necessarily mean that these are true edges. These diagrams thus provide an assessment of the concordance of the different inference methods.

Other noteworthy trends are the large overlap between PIDC, CLR and PUC; more surprising perhaps is the apparent similarity of the signal picked up by the two correlation methods and MI (Fig. 2). Furthermore, GENIE3 appears to be an outlier and routinely scores a relatively sizeable set of candidate edges that are not picked up by any other method. In the absence of a ground truth it is hard to make too much of these Venn diagrams, except perhaps at the extremes: groups of strong methods are expected to result in high concordance (reflected in large overlap), whereas very small overlap may indicate a set of three particularly poor inference methods.

### 3.4 Behaviour under artificial noise

In order to investigate how sensitive functional assortativity is to the assumption of mixing patterns, we show in Fig. 3 its behaviour as the inferred networks are perturbed in different ways.

**Figure 3:**
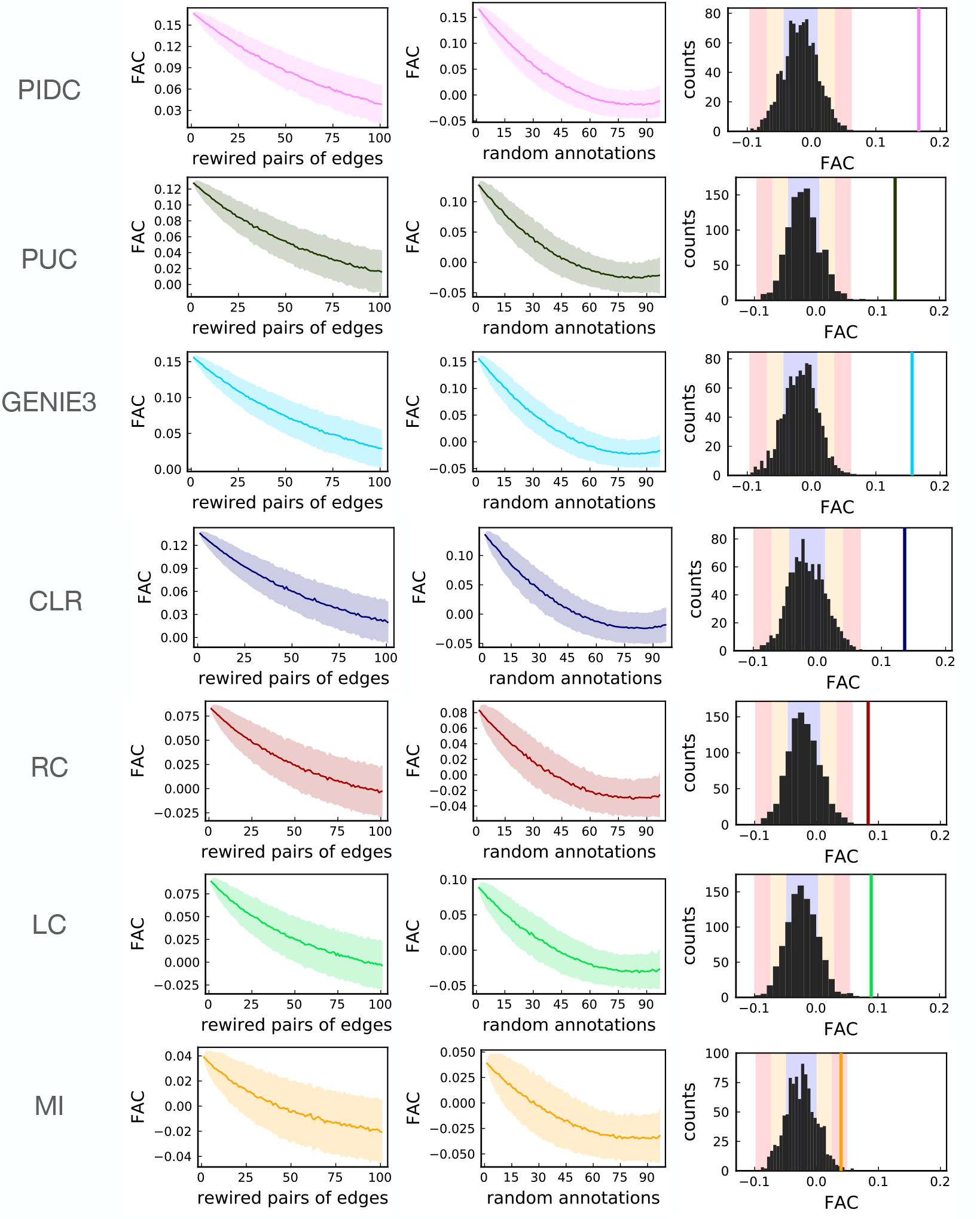
Illustrating the behaviour of the FAC under noisy conditions. Mean (solid line) and standard deviation from the mean (shaded area) of the FAC as pairs of edges are rewired at random (left column), and as nodes are randomly attributed a different annotation (central column) – each plot shows 1000 repeats. Right column: comparison of the observed FAC for networks with 200 edges (vertical line) against distributions of the FAC in 1000 random networks with 200 nodes; blue, orange, and red coloured bands respectively indicate one, two and three standard deviations from the mean.

We find that the FAC tends to 0 as biological functions are randomised among the nodes (Fig. 3A), showing that the signal it picks up is not merely an artefact of a particular network topology. Instead this suggests that the inferred networks pick up a real signal from the nodes, which is a non random function of the particular topology of inferred networks and the associated group labels.

This is supported by the signal disappearing into noise with increasing levels of randomness in network structure (Fig. 3B) and a sanity check of random values as expected in random networks (Fig. 3C).

From this, we conclude that functional assortativity is *informative* and *reliable*. Informative, because it is different than random: it measures the extent of mixing patterns by function, and the values it takes are not the result of chance alone. Reliable, because it is robust to low levels of noise – it can still pick up a signal under reasonable perturbations – but that signal vanishes for higher levels of noise, thus apparently avoiding false positives.

## 4 Discussion

The lack of comprehensive, experimentally-derived networks that can be used as a reference makes rigorous assessment of network inference algorithms challenging. Most methods have their specific assumptions and this will lead to discrepancies in their predictions.

In the context of analysing real biological networks, such discrepancies are a clear indication that rankings of network inference algorithms should be taken with caution: they are only a reflection of their performance in the specific context they were tested in (and indeed, for the same inference method, we have seen discrepancies in performance – e.g. excellent predictions in some contexts, but only slightly better than random in others [21]). This goes to show that there is no definitive “best” method and performance is context-dependent.

We argue that this motivates the need for ways to compare inferred networks that are not biased towards our necessarily limited current knowledge [39]. We believe that the assumption of mixing patterns by function achieves this: it uses *expectations* as a basis for comparison, and these expectations are backed by both theoretical arguments and empirical results. This frees us of the potentially misleading circularity that is inherent to *in silico* approaches, and has the advantage of making our assumptions explicit and thus falsifiable.

We find that the behaviour of mixing patterns by function is reliably measured by the FAC. This makes it conceptually related to network modularity, where instead of quantifying aspects of network structure based purely on topological properties, it does so based on biological function. This balances the limited mechanistic assumptions of many network inference methods (although GENIE3 and other methods allow inclusion of prior knowledge) – only quantifying statistical dependency at its core – by grounding the process in realistic biological assumptions.

While clearly not all interactions are between genes performing the same biological function, this type of interaction will dominate (compared to the case of purely random connections). Thus functional assortativity allows us to quantify confidence in inferred networks as we would thus put more trust in networks that are functionally assortative than those that are not. As such, it is a heuristic that can guide the decision-making part of the inference process when it is understood as an inverse problem [40]. It effectively displaces the notion of confidence from the ability to reproduce previous observations to ability to produce expected results. We believe this approach, and others based on a similar perspective, to be useful in contexts where our knowledge is limited.

## 5 Conclusion

Networks remain a useful starting point for mechanistic analysis and assessing confidence in *in silico* inferred networks is important for the further use of such networks. Two limiting factors in our approach are (i) it only provides a heuristic way of ranking different inferred networks; and (ii) it requires that genes be annotated with a biological function [41, 42] – this data may not be readily available; it may be incomplete; and it may be subject to uncertainty and or errors.

The present approach relies on the annotation of nodes, and increasing the quality of such annotations will clearly benefit this proxy measure. Additional improvements could come from considering functional assortativity locally, that is in specific areas of the overall network.

Currently, however, as a rule of thumb, functional assortativity allows us to rank different candidate networks or network inference methods. knowing which inferred networks are worth further consideration, and which ones are best ignored will have a profound impact on our ability to make use of networks. Quickly being able to reject some network inferences does allow for more streamlined analysis, but is also essential [43] if we want to base predictions on ensembles of network inference methods.

We believe that there is an urgent need for an approach such as the one described here. In the absence of rigorous statistical assessments of inferred networks, the simple heuristic provided by the functional assortativity coefficient can provide criteria by which to gauge the reliability of inferred networks.

## Data and methods availability

All data and code are available at http://doi.org/10.5281/zenodo.4021679.

## Acknowledgments

We gratefully acknowledge the help of Thalia Chan during the early stages of this research. We thank the members of the *Theoretical Systems Biology Group* at Imperial College London and the University of Melbourne for many helpful discussions. This work is funded through the University of Melbourne’s Deputy Vice Chancellor of Research Fund.

